# Habitat filtering differentially modulates phylogenetic vs functional diversity relationships between dominant ground-dwelling arthropods in salt marshes

**DOI:** 10.1101/2020.06.19.161588

**Authors:** Aurélien Ridel, Denis Lafage, Pierre Devogel, Thomas Lacoue-Labarthe, Julien Pétillon

**Affiliations:** UMR CNRS 6553 Ecobio, Université de Rennes, 263 Avenue du Gal Leclerc, CS 74205, 35042 Rennes cedex, France; UMR CNRS 7266 LIENSs, Université de La Rochelle, 2 Rue Olympe de Gouges, 17000 La Rochelle, France

**Keywords:** spiders, carabids, salinity, NW France, species richness, traits

## Abstract

While mechanisms underlying biological diversities at different scales received huge attention over the last decades, whether local abiotic factors driving functional and phylogenetic diversities can differ among ecologically and phylogenetically closely related taxa remains under-investigated. In this study, we compared correlations and drivers of functional (FD) and phylogenetic (PD) diversities between two dominant taxa of ground-dwelling arthropods in salt marshes, spiders and carabids. Pitfall trapping in two sampling sites of N-W France resulted in the collection and identification of more than 7000 individuals belonging to 67 species. Morphological and behavioral traits, as well as molecular sequences of COI gene, were attributed to all species for calculating functional and phylogenetic diversities respectively. Both taxa exhibited high correlation between FD and PD, which was even higher in carabids probably due to their lower species richness. Analyses using Bayesian framework and structural equation modeling revealed that FD and PD were positively influenced by taxonomic diversity in spiders and carabids, but abiotic factors driving FD and PD differed between taxa. Salinity especially drove the taxonomic diversity of carabids, but not that of spiders, suggesting that spiders are more plastic and less selected by this factor. Phylogenetic diversity was conversely influenced by salinity in spiders but not in carabids. This interesting result can be interpreted by different evolutionary history and colonization process of salt marshes between the two model taxa. Our study finally highlights that, even in taxa of the same phylum and occupying the same niche in a highly constrained habitat, functional and phylogenetic diversities can have different drivers, showing different filtering mechanisms and evolutionary history at small spatial and temporal scales.

## Introduction

Describing spatial patterns of species assemblages is an objective of community ecology that can be directly used for biological conservation (Devictor *et al*., 2010; Arnan *et al*., 2016). The study of factors driving local diversity is an essential step to understanding these patterns, and has long been performed using taxonomic diversity only. However, this approach does not consider all facets of biodiversity such as accumulated evolutionary history traits (Webb *et al*., 2002), or the diversity of morphological, physiological and ecological traits of an assemblage (Petchey and Gaston, 2002, 2006). Taking into account both Taxonomic (TD), Phylogenetic (PD) and Functional (FD) Diversities appear to be essential for better understanding composition and dynamics of species assemblages (Webb *et al*., 2002). In the same way, these metrics are seen as allowing a better assessment of biodiversity in conservation studies, because they are the three major components of biodiversity (Swenson, 2011). In a complementary way, TD can bring information about species composition of an ecosystem resulting from several processes, while PD highlights a part of these process by providing information on the evolutionary relationships among coexisting species (Webb *et al*., 2002). On the other hand, FD reflects links between biodiversity, ecosytemic functions and environmental constraints (Mouchet *et al*., 2010), such as functional response of species assemblages to environmental filtering (Diaz *et al*., 2007).

Despite the fact that these metrics are seen as complementary, their mutual relationships remain unclear (Devictor *et al*., 2010). Instinctively, a positive correlation between TD and PD or FD is expected because the presence of more species can indirectly capture more lineage and functional traits. However, it has been shown that a similar number of species can be associated with different levels of PD and/or FD (Safi *et al*., 2011; Tucker and Cadotte, 2013). Moreover, the strength of the correlation between TD and PD likely depends on the number of accumulated evolutionary history traits, but can also be influenced by other parameter like the symmetry of the phylogenetic trees, branches length, species pool size and spatial autocorrelation (Tucker and Cadotte, 2013). Additionally, PD is often seen as a proxy of FD because functional traits of an assemblage indirectly reflect evolutionary history of this assemblage (Winter *et al*., 2013). If traits are phylogenetically conserved, PD can provide also an information about unmeasured functional traits (Cadotte *et al*., 2009). In contrast, some studies revealed a fluctuating relationship between PD and FD (Devictor *et al*., 2010; Cadotte and Tucker, 2018), which may depend on the studied species pool size (Mazel *et al*., 2018). In addition to that, the relationship between PD and FD can also depend on the shape of phylogenetic tree (Letten and Cornwell, 2015), as well as on the number of used functional traits (Tucker *et al*., 2018). Finally, taking into account taxonomic diversity in both PD and FD can lead to a correlation between them as a side effect (Safi *et al*., 2011), because both of PD and FD include TD for their calculations (Pavoine *et al*., 2013; Cadotte *et al*., 2019). To better understand the relationships between diversity metrics, it is essential to understand what are the drivers of these metrics, and how they affect the relationships between these metrics. If the drivers of TD have been studied for a long time, disentangling the influence of factors driving PD and FD is a more recent challenge (Fournier *et al*., 2017; Hanz *et al*., 2018; Fichaux *et al*., 2019). The number of studies dealing with the taxonomic, phylogenetic and functional facets of diversity concomitantly is still too small (Cadotte *et al*., 2019). However, it seems essential to carry out research with an overview of diversity components to highlight their causes and consequences for ecosystem functioning (Fournier *et al*., 2017). In addition, there is a lack of knowledge about PD and FD drivers in highly constrained environments (Cadotte *et al*., 2019), and particularly on taxa occupying similar ecological niches at a small spatial scale.

Here we propose to carry out a multi-taxa approach taking into account all facets of biodiversity in salt marshes, a highly constraint environment. Salt marshes are transitional ecosystems between marine and terrestrial systems (Adam, 1990). Due to their intertidal position, salt marshes are subject to several environmental stress, including periodic flooding and the resulting salinity gradient. These stress have a strong impact on salt-marshes’ organisms (Lefeuvre *et al*., 2003), and most of the species occuring in these ecosystems have a high phenotypic plasticity, or even morphological, physiological or behavioural adaptations to cope with these stresses (Pennings & Callaway, 1992; Pétillon *et al*., 2009, 2011). Among these organisms, terrestrial arthropods constitute the most diverse and abundant group in salt marshes (Desender & Maelfait, 1999; Pétillon *et al*., 2014), and especially spiders and carabids for predatory arthropods (Ford *et al*., 2013; Coccia and Fariña, 2019). In addition to that, these two taxa play important functional roles in these environments where they act as both prey and predators (for carabids and spiders, see Raino & Niemelä., 2003 and Nyffeler & Birkhofer, 2017, respectively). For spiders and carabids, how environmental variables and taxonomic diversity influence relationship between PD and FD remain unknown in this type of harsh ecosystem. In this study, we investigated the correlation between PD and FD, allowing the simultaneous consideration of diversity metrics as well as environmental factors. More precisely, we tested the following hypotheses to better understand the mechanisms shaping diversities of arthropods in salt marshes:

### Hypothesis 1

in a gradually constraining environment such as salt-marshes, we expect a strong correlation between phylogenetic and functional diversity for both taxa studied (Cadotte *et al*., 2009; Winter *et al*., 2013), because abiotic filters select for a small number of specialized species adapted to high soil salinities and frequent inundations (Foster & Trehern, 1976; Foster, 2000). In addition, this relationship is expected to be stronger in carabids because of the higher phylogenetic proximity of species with conserved and functionally adapted traits to salt marshes carabids (Desender & Serrano, 1999; Van Belleghem *et al*., 2015), compared to spiders (Pétillon *et al*., 2011; Puzin *et al*., 2014).

### Hypothesis 2

we expect that TD influence the strength of this correlation between PD and FD by side effect (Safi *et al*., 2011; Pavoine *et al*., 2013; Cadotte *et al*., 2019). Due to the greater sensitivity of carabids to environmental constraints such as salinity (Pétillon *et al*., 2008), the relationship between TD and both PD and FD diversity is expected to be stronger for this taxa, resulting in a more specialized pool of species.

### Hypothesis 3

The environmental variables are expected to influence taxonomic diversity in the same way for each taxon. More precisely, salinity is expected to have a negative influence on taxonomic diversity of both spiders and carabids because it acts as a highly constraining factor for terrestrial species (Pétillon *et al*., 2008). In addition, the environmental factors should influence similarly the PD of salt-marshes organisms, whatever the taxon, because of their recent evolutionary history in salt marshes. In contrast, environmental variables affecting functional diversity are also expected to differ between taxa as shown in other ecosystems (Fournier *et al*., 2015).

## Material and methods

### Study sites and sampling design

The study was conducted on two salt marshes in Charente-Maritime (New Aquitaine Region, France). The First site (site 1) is located on 46.22885, -1.50112; and the second site (site 2) on 45.88503, - 1.094999 (Fig.1-A). This study focuses on the part of each of these sites classified as a national nature reserve, and were selected because they have a large surface area of coastal salt marshes (EUNIS A2.5), habitats considered to be of high heritage value and targeted in this study.

**Figure 1.**
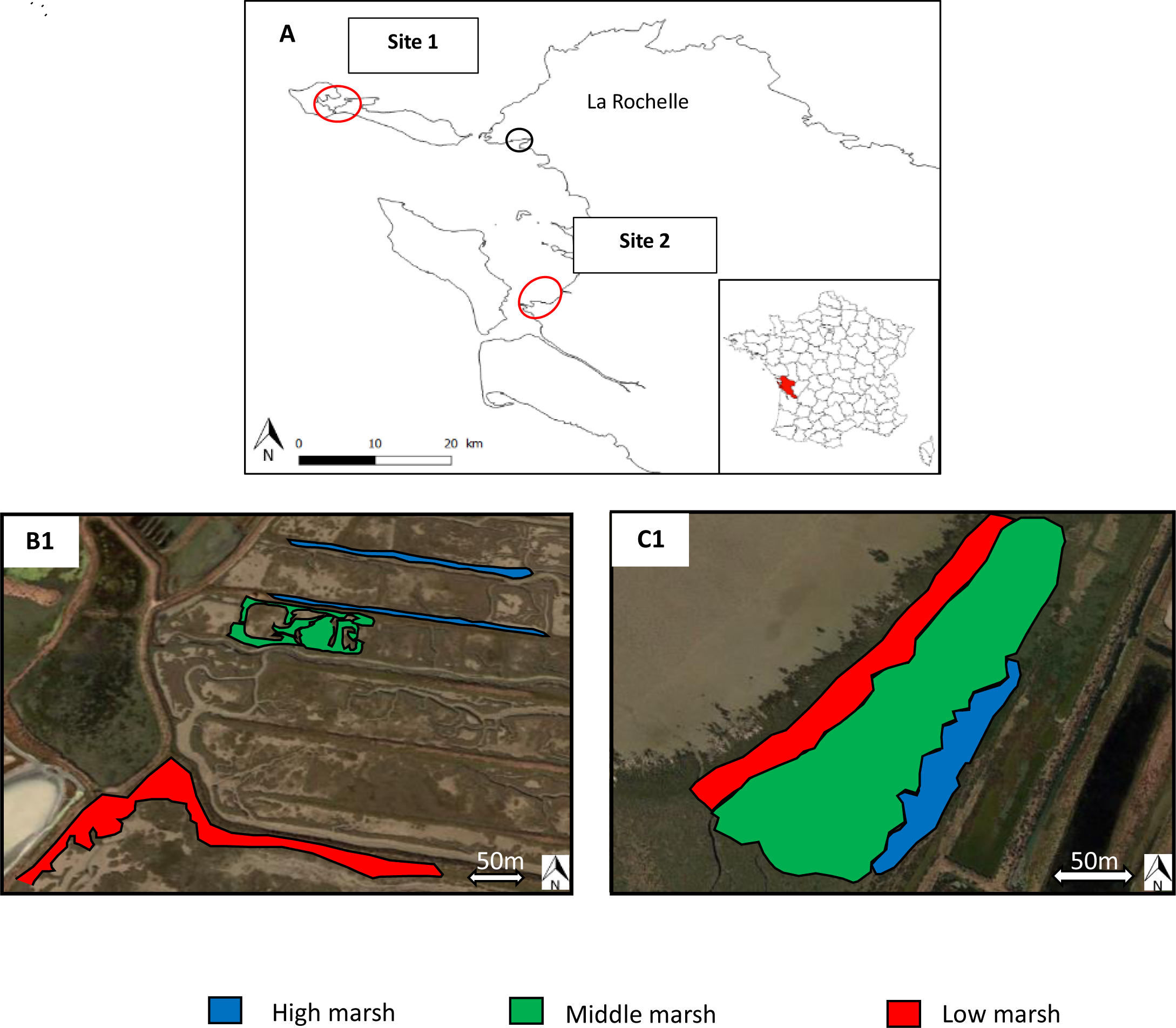
Location of the study sites within the Charente-Maritime department (Western France). (A) Site one located on the Ile de Ré, and the second site located in the municipalities of Moëze and Saint-Froult. Location of the three salt-marsh zones defined on the basis of plant composition / species on the first site (B1) and the second site (C1) HM=high marsh (blue); MM=middle marsh (green); LM=low marsh (red).

Three zones have been defined per site according to their topographic location and vegetation characteristics (successively: high, middle and low marsh), using EUNIS typology. These habitats are representative of different immersion frequencies and the resulting salinity gradient present on each site. Therefore, identified zones within which immersion frequencies and vegetation are considered homogeneous. On site 1, the zones are distant and fragmented as a consequence of past salt farm on this site (Appendix 1-B). The high marshes have about 0,5% of annual recovering tides are whereas the lowest areas have about 60%. On site 2, the zones are continuous from the pioneering to the upper salt marshes (Appendix 1-C). The high marshes have about 10% of annual recovering tides and the low marshes have about 35%.

Pitfall trap, consisting in cylindrical PVC pipes (diameter of 9 cm) and buried in the ground so as to form a continuity between the upper edges of the pipe and the ground, were used to sample ground-active spiders and carabids (Curtis, 1980). Inside this pipe, a plastic jar containing a preservative liquid is inserted. Over this container, a funnel attached to the outer edges of the PVC tube was placed. Finally, a roof was set over the device supported by metal stakes, to protect the collecting liquid from a water supply.

The collecting plastic jar were filled at 3/4 with a 250 g.L^-1^ saline solution supplemented by a drop of dishwashing liquid per litre (which reduces the surface tension of the liquid). Each of the study sites therefore has three zones, each with a total of three sampling stations to provide sufficient spatial replication. Each station includes four pitfall traps arranged linearly. The interception radius of a pitfall trap being of around 5m (Topping & Sunderland, 1992), traps were placed in the centre of non-overlapping circles of 10m in diameter, in order to prevent them from influencing each other. A total of 36 traps were set up per study site. Five trapping sessions of 4 to 12 days each were carried out between April and July 2019, each comport between 4 and 12 days per site. The dates and duration of the sessions differ between sites due to their different immersion frequencies, resulting in 29 days of continuous sampling carried out on the first site, and 48 days of continuous sampling for the second site. Carabids and spider species have been identified to the finest possible taxonomic level.

### Environmental variables

When setting pitfall traps, environmental variables were recorded within a 5m radius circle around each pitfall trap (matching to the theoretical the interception areas of the traps). Litter depth and average and maximum height of vegetation were measured to the nearest centimetre, the percentage of bare ground estimated by eye to the nearest, and the soil salinity was estimated by measuring the conductivity of one gram of soil diluted in 15 ml of distilled water. Within the same areas, phytosociological survey (Braun-Blanquet, 1928) were carried out with an allocation of a recovery percentage to the plant species present, in order to check whether the zones are homogeneous in terms of vegetation. (Table 1-B).

**Table 1.**
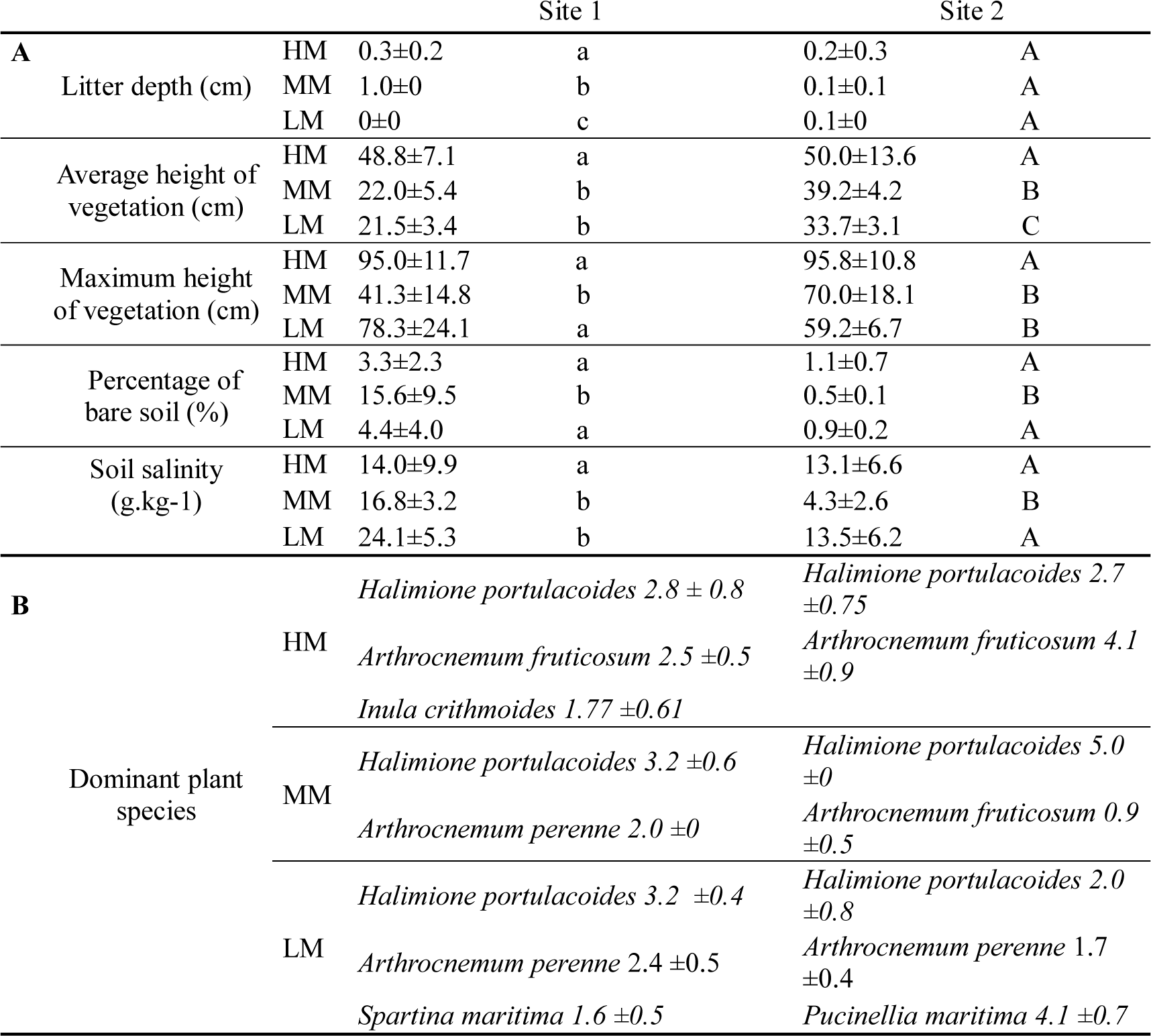
Environmental variables (mean ± sd, N=12) for each salt-marsh zone and for each site Successive letters indicate significant differences by Anova followed by Tukey post-hoc tests of or by Kruskal-Wallis followed by Mann-Whitney or Welch tests when appropriate. A Bonferroni correction was used for post-hoc tests. Plant species are given only when they occurred in more than 75% of the surveys carried out in the area and had a coverage >1.5).

Litter depth and vegetation height (average and maximum) tend to decrease from the high to the low marsh, with some differences noted for the site one (Table 1-B). The percentage of bare ground is maximum in the middle marshes for the site 1, and minimal in this same area for the site 2. Salinity is globally increasing from high to the low marsh, although this variable is minimal in middle marshes for the site 2. According to phytosociological survey, the low marshes can be characterized by a dominance of *Halimione portulacoides* for the site 1 against a dominance of *Pucinellia maritima* in the site 2. The middle marshes are dominated by *H. portulacoides* whatever the site. Finally, a co-dominance of *H. portulacoides* and of *Arthrocnemum fruticosum* was found on the site 1, contrasting to a dominance of *A. fruticosum* on the site 2.

### Phylogenetic tree building

Phylogenetic trees were constructed by combining phylogenetic and taxonomic data from literature, assuming identical branch length between genus (1) and species (0.5) as distances were not available for all species and we wanted to be able to compare results between taxa. The spiders phylogenetic tree (Appendix 4) was adapted from Wheeler *et al*., (2017). Genus that were not present in Wheeler’s tree were placed using Arnedo *et al*., (2009), Frick et al. (2010), Wang et al. (2015) for Linyphiidae and Agnarsson, (2004), Azevedo et al. (2018), Maddison (2015), Millidge (1977), Piacentini & Ramírez (2019) and Scharff et al. (2019) for other families. The carabids phylogenetic tree (Appendix 5 was adapted from López-López & Vogler (2017), Martínez-Navarro, Galián, & Serrano (2005), Ober & Maddison (2008), Ruiz, Jordal, & Serrano (2009) and Sasakawa & Kubota (2007)

### Functional traits used

In order to calculate the functional diversity per pitfall trap for each taxon, functional traits were assigned to each of the spider and carabids species according to the literature (Appendix 1). Selected traits were: size, dispersal capacity and overall diet of species of each group.

## Statistical analysis

Taxonomic diversity was estimated by the species richness of samples, computed with the BAT package (Cardoso *et al*. 2015) that uses the proportion of singleton or single species and following Lopez *et al*. (2012) methods to obtain corrected jackknifes estimates. Final taxonomic diversity was calculated by averaging corrected Jackknifes estimators in order to account for sampling variability. In the same way, phylogenetic and functional diversities were estimated following Cardoso *et al*. (2014) methods using BAT package. Final PD and FD were also calculated by averaging corrected Jackknifes estimators. Distance matrix for phylogenetic distances was calculated with gower distance from the FD package (Laliberté *et al*., 2014).

The correlation between PD and FD was estimated in a Bayesian framework with a Student’s t distribution (which reduces sensitivity to outlayers) with the brms package (Bürkner, 2018). We used 2000 iterations on 4 chains. Model convergence was check by visually inspecting diagnostic plots. To select environmental variables affecting PD and FD (for later use in Structural Equation Models), models were built within a Bayesian framework using brms (Bürkner, 2018) with two chains and default priors. All environmental variables were standardized and centered. The models included salinity, % bare-ground, litter depth, mean vegetation height, maximum vegetation height and salt-marsh zone. Model convergence was checked by visually inspecting diagnostic plots and using Rhat value. Parameter selection was based on “HDI+ROPE decision rule” (Kruschke & Liddell, 2018) with a range value determined as -0.1 * sd(y), 0.1 * sd(y) (Kruschke & Liddell, 2018) and was performed using bayestestR (Makowski *et al*., 2019).

We assessed the relative contribution of environmental variables selected by Bayesian models using structural equation modeling (SEM). The SEM approach also allows us to assess the indirect effect of TD on PD and FD as their calculation both depends on TD and to test for correlated errors between PD and FD. A significant correlated error between the two variables would indicate the existence of an unknown parameter influencing both variables. We used the piecewise SEM package (Lefcheck, 2016) as it allows us to use mixed models in association with nlme package (Pinheiro *et al*., 2019). Our initial model included the following links: (1) PD is affected by TD and selected environmental variables, (2) FD is affected by TD, selected environmental variables and PD (3) TD is affected by selected environmental variables and (4) there is correlated error between PD and FD. Site was used as a random factor in every link modeled using nlme (Pinheiro *et al*., 2019). After the specification of the initial model, we re-defined our model excluding non-significant links (p<0.05) using a stepwise approach until ΔAICc <2 between two subsequent models. Finally, we assessed model fit using Fisher’s C statistic. All statistical analyses were performed with R Studio software (version 3.5.1).

## Results

A total of 3359 adult spiders belonging to 55 species, of which 58.9% of individuals are considered halophilic (Appendix 2), was collected by pitfall traps (Ridel *et al*., 2020). Spiders had an average size of 7.13 ± 3.79 mm. Hunting guilds were dominated by ground-hunting individuals (56.6%), and for dispersal methods, most individuals were ballooners (63.4%).

A total of 4005 carabids belonging to 12 species, of which 99.7% of individuals sampled are considered halophilic (Appendix 3), was collected by pitfall traps. Carabids were 6.82 ± 0.84 mm sized on average. Diet of carabids was dominated by generalist predator individuals (99.3%), and the main dispersal technique is represented by polymorph individuals (92.5%).

The correlation factors between PD and FD was 0.48 (95% CI: 0.27-0.66) and 0.89 (95% CI: 0.83-0.94), for spiders and carabids, respectively (Fig. 2).

**Figure 2.**
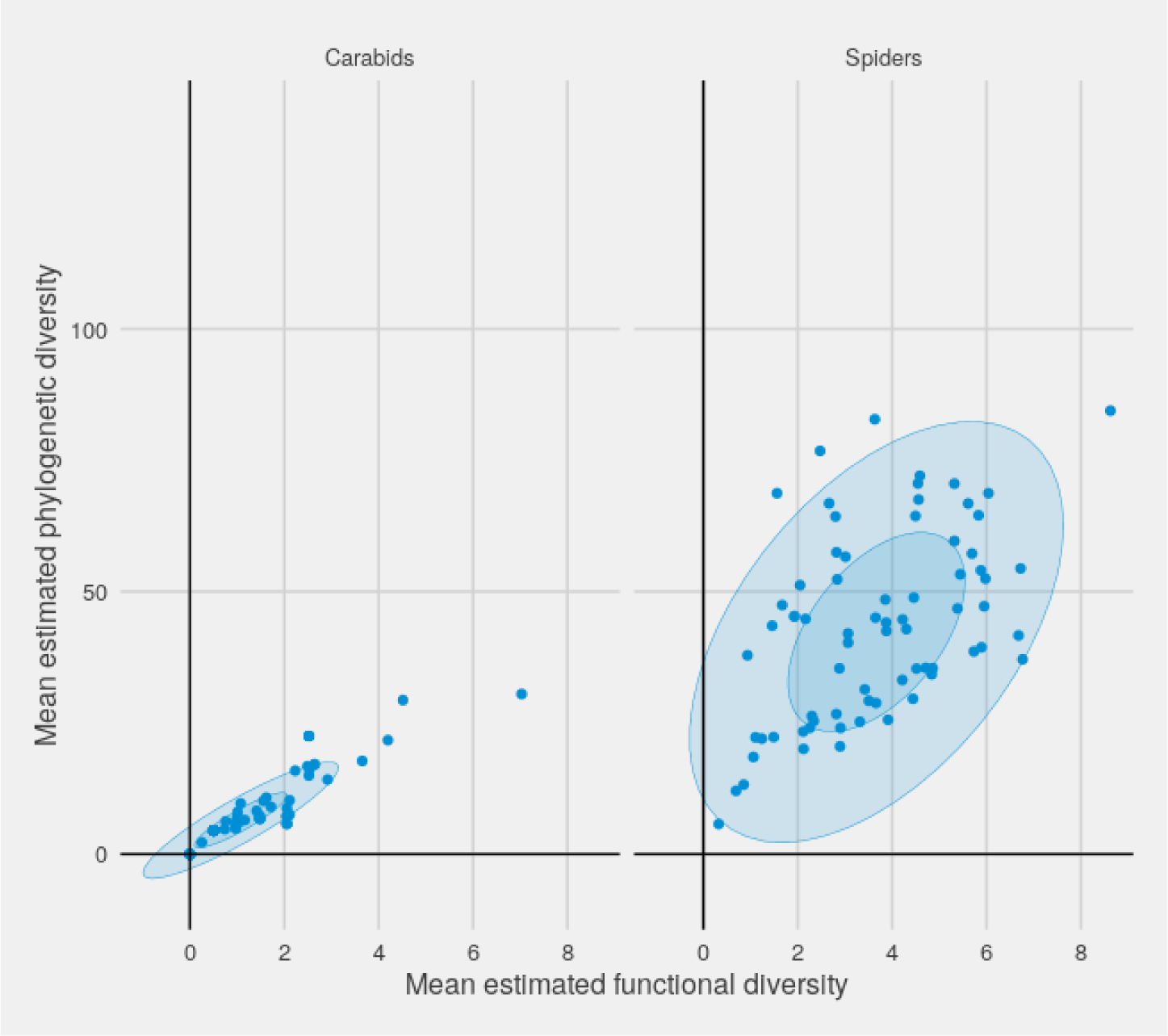
Plot of mean phylogenetic diversity as a function of functional diversity for spiders and carabids. Ellipses correspond to 95% confidence intervals.

The Bayesian model for spider PD successfully converged, and had a R^2^ = 0.271. Mean vegetation height and salinity were the best explanatory variables. Mean vegetation height effect on spider PD had a high probability of existing (pd = 98.65%, Median = 8.49, 89% CI [2.70,14.37]), and could be considered as significant (0% in ROPE). Salinity effect on spider PD had a high probability of existing (pd = 99.7%, Median = 7.14, 89% CI [3.57, 11.11]), and could be considered as significant (0% in ROPE). The model for spider FD successfully converged, and had a R^2^ = 0.415. Litter depth was the best explanatory variable. Litter depth effect on spider FD had a high probability of existing (pd = 99.6%, Median = -0.86, 89% CI [-1.31, -0.35]), and could be considered as significant (0% in ROPE). The model for spider TD successfully converged, and had a R^2^ = 0.442. Mean vegetation height was the best explanatory variable. Mean vegetation height effect on spider TD had a high probability of existing (pd = 99.9%, Median = 4.76, 89% CI [2.21, 7.29]), and could be considered as significant (0% in ROPE).

The Bayesian model for carabids PD successfully converged, and had a R^2^ = 0.188. Mean vegetation height and salinity were the best explanatory variables. Mean vegetation height effect on carabid PD had a medium probability of existing (pd = 97.75%, Median = -0.58, 89% CI [-0.99, -0.11]), and its significativity remained undecided (7.58% in ROPE). Salinity effect on carabid PD had a medium probability of existing (pd = 95.75%, Median = -1.79, 89% CI [-3.33, 0.31]), and its significativity remained undecided (7.00% in ROPE). The model for carabids FD successfully converged, and had a R^2^ = 0.18. Mean vegetation height was the best explanatory variable. Mean vegetation height effect on carabid FD had a high probability of existing (pd = 99.6%, Median = -0.86, 89% CI [-1.31, -0.35]), but its significativity remained undecided. The model for carabids TD successfully converged, and had a R^2^ = 0.222. Mean vegetation height and salinity were the best explanatory variables. Mean vegetation height effect on carabid TD had a high probability of existing (pd = 99.15%, Median = - 1.81, 89% CI [-2.92, -0.71]), and could be considered as significant (0% in ROPE). Salinity effect on carabid TD had a probability of existing (pd= 97.70%, Median = -0.89, 89% CI [-1.552, -0.219]), still its significativity remained undecided (2.9% in ROPE).

When testing the relationships between the different metrics of diversity and environmental variables for spiders, our final SEMs indicated good fit with the data (Fisher’s C = 1.075, p = 0.898; Fig. 3). Salinity and litter depth were linked respectively to spider PD (linked selected in the model but only marginally significant: p =0.068) and FD (linked selected in the model and again almost significant: p =0.051). Spider PD was strongly and positively related to TD (coefficient standard estimate: 0.705). Spider FD was positively linked to PD (coefficient standard estimate: 0.231) and TD (coefficient standard estimate: 0.435).

**Figure 3:**
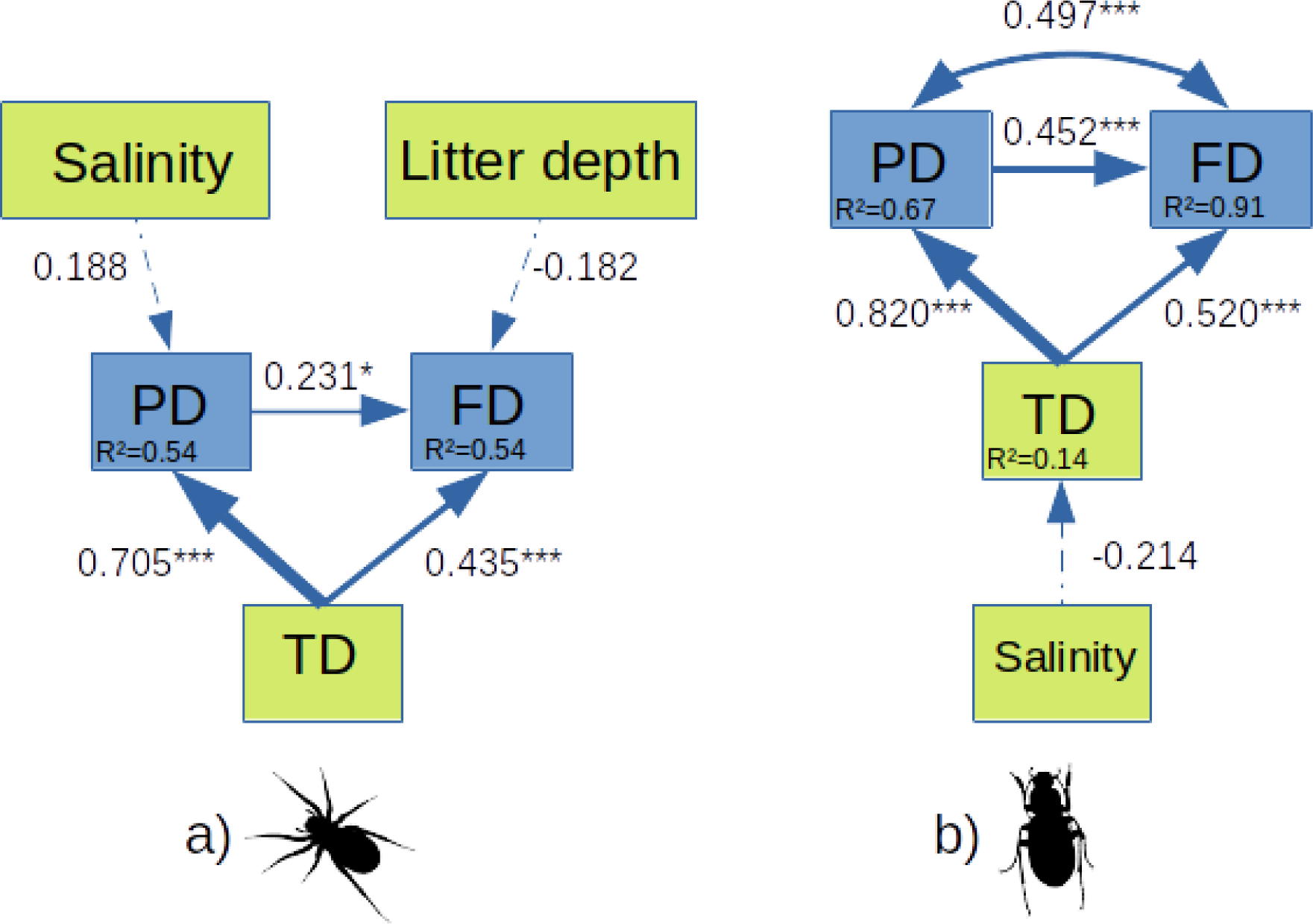
Best piecewise SEMs showing links between a-, phylogenetic and functional diversity and environmental variables for a) spiders and b) carabids. Thickness of arrows is proportional to the standardized path coefficients (directionality and size given within boxes). Asterisks give significance level of linkages (<0.1, *<0.05, **<0.01, ***<0.001), and dashed lines correspond to paths included but not significant (p>0.05). Double arrow represent correlated errors. Conditional R^2^ values are given within the boxes containing variables.

When testing the relationships between the different metrics of diversity and environmental variables for carabids, our final SEMs indicated good fit with the data (Fisher’s C = 5.368, p = 0.252; Fig. 3). Salinity was negatively linked to carabid TD (linked selected in the model but not significant: p =0.110). Carabids PD was strongly and positively related to TD (coefficient standard estimate: 0.821). Carabids FD was positively linked to PD (coefficient standard estimate: 0.452) and TD (coefficient standard estimate: 0.520). Correlated errors were found between FD and PD (coefficient standard estimate: 0.497).

## Discussion

### Correlations between phylogenetic and functional diversities

As expected, positive correlations between PD and FD diversity metrics were found for both spiders and carabids. This result can be explained by the fact that in highly constrained habitats such as salt marshes, environmental filtering leads to a selection of species according to functional groups (Ford *et al*., 2013), which is the case here for species from both taxa. In this case, the positive correlation between PD and FD are likely due to the fact that functionally adapted species are also often phylogenetically close (Winter *et al*., 2013). Interestingly, this link between PD and FD is expected to increase with the number of functional traits used (Tucker *et al*., 2018), but here only three functional traits were used to homogenize their number and nature between the two studied taxa. This relationship is strong here even with a small number of traits used, which suggests that in salt marshes functional traits are phylogenetically conserved. This interesting hypothesis should still be tested further, which was already stressed by Cadotte et al. (2009). Also, the small number of species encountered in studied salt-marshes (N=55 for spiders and N=12 for carabids) compared to less constrained environments from the same biogeographic area (see e.g. Lafage et al. 2015: N=99 for spiders and N=43 for carabids) may also increase the strength of the PD-FD correlation (Tucker and Cadotte 2013). This result comforts the idea that salt marches present strong environmental constraints that act as strong filter, selecting functionally adapted species to flood and salinity from a larger regional species pool (Pétillon *et al*., 2008, 2014).

The observed links between phylogenetic and functional diversities can also result from parameters of phylogenetic trees used. There is evidence that the strength of PD-FD correlation can be increased by the use of phylogenetic trees that are symmetrical and/or have long terminal branches (Tucker and Cadotte 2013). Here, it was unfortunately not possible to calculate an index of symmetry and/or length of the terminal branches because branch had similar length between genus and species due to limited knowledge in the phylogeny of arthropods. Finally, it is also possible that the inclusion of taxonomic diversity in the calculation of phylogenetic and functional diversities influences the relationship between these two metrics (Safi *et al*., 2011; Pavoine *et al*., 2013: see discussion below).

The correlation between PD and FD was stronger for carabids than for spiders, also following our initial expectation. First, the reduce number of salt-marshes occurring carabids compared to spiders (i.e. 12 species for carabids against 55 species for spiders) increase this correlation (Tucker and Cadotte 2013). Moreover, species of carabids adapted to salt marshes are phylogenetically closer than spiders ones due to a more recent evolutionary radiation of this taxon in salt marshes (see Desender & Serrano, 1999 and Van Belleghem *et al*., 2015). In fact, it should be noted that the species of halophilic spiders are not concentrated in the same genus (Pétillon *et al*., 2011; Puzin *et al*., 2014), but rather result from adaptation of particular species independently of their phylogeny (traits convergence: Spicer & Gaston 1999). Therefore, the observed relationship between phylogenetic and functional diversities suggests that functional traits are phylogenetically more conserved in carabids.

### Effects of taxonomic diversity on PD-FD relationships

First, the SEMs carried out on both taxa revealed that the links between phylogenetic and functional diversity was weaker than the links between taxonomic diversity and both of phylogenetic and functional diversities (see Fig.3). These results suggest that the relationship between phylogenetic and functional diversity is mainly affected by the taxonomic diversity, agreeing with our initial expectations and previous studies (Safi *et al*., 2011; Pavoine *et al*., 2013; Cadotte *et al*., 2019).

Unsurprisingly, taxonomic diversity was also found to be more strongly related to the other diversity metrics considered in carabids than in spiders. One explanation could be that carabids are more sensitive to salinity than spiders because of their ecology (larval stages on or in the soil, adults less in the vegetation than e.g. spiders; Luff 2007). So, carabid species of salt marshes are more specialised to this environment than spiders, and indeed possess spectacular morphological adaptations like a waterproofing inter-tegument cuticle (Foster & Treherne, 1976), which altogether results in a restricted pool of species mainly composed of halophilic species. Furthermore, as stated above, the halophilic carabids appear to be more phylogenetically clustered than halophilic spiders, resulting in a strong link between taxonomic diversity and both phylogenetic and functional diversities for this taxa. The lower percentage of halophilic spider species found in this study compared to that of halophilic carabids is consistent with this hypothesis (respectively 58.9% vs 99.7%).

### Effects of environmental filtering on diversities metrics

Surprisingly, the salinity influenced the taxonomic diversity of carabids – because of the their greater sensitivity to this stress (see above and Irmler *et al*., 2002; Pétillon *et al*., 2008) - but not spiders taxonomic diversity, whereas the importance of this factor in structuring spiders assemblages was reported from previous field studies (Pétillon *et al*., 2005, 2008). This unexpected results could be explained by the fact that spiders are more plastic to saline stress than carabids (Pétillon *et al*., 2008, 2011), and thus that more diverse non-specialized spiders could use salt marsh habitats. Laboratory experiments indeed shown repeatedly that halophilic spiders, although strictly restricted to salt marshes, actually performed better (in terms of both survival and fitness) without saline stress than under saline to hyper saline conditions (see: Foucreau et al. 2012, Renault et al. 2016). For carabids, salinity likely had a strong influence on this taxon because of their greater sensitivity to this stress (see above and Irmler *et al*., 2002; Pétillon *et al*., 2008).

Interestingly, the phylogenetic diversity is differentially influenced by the salinity for spiders and by an unidentified variable for carabids. Again, these results are consistent with the hypothesis of a stronger plasticity of spiders in response to salinity (Pétillon *et al*., 2008, 2011). This higher plasticity could indeed result in a less specialized pool of spider species as suggested by low percentage of halophilic species recorded in traps. On the other hand, since environmental filters act more strongly on carabids, it seems logical that salinity had little influence on their phylogenetic diversity because it is composed by a pool of species already strongly selected by this factor.

Conforming our initial expectation, the main environmental variable driving the functional diversity of carabids and spiders differed between these taxa, with an influence of litter depth for spiders and an unidentified variable for carabids. These results are in line with the idea that the drivers acting on the functional diversity of these taxa can be different (Schirmel *et al*., 2012; Schirmel and Buchholz, 2013; Fournier *et al*., 2015; Schirmel *et al*., 2016). The effect of litter on the functional diversity of spider is consistent with Uetz (1979) and Döbel et al. (1990). On the other hand, it is important to point out that the unidentified environmental variable affecting the functional diversity of carabids is the same that the one driving their phylogenetic diversity, thus strengthening again the links between these two metrics. Parameters, not measured in this study, such as soil moisture and the duration of flooding by the adjacent sea are known to affect the functional diversity of carabids (Fournier *et al*., 2015), and can certainly be candidates for this unidentified variable. The sediment granulometry is also known as functional diversity filter by driving the carabids endogenous larval life and the full-grown burrowing-type strategy (Antvogel and Bonn 2001). The SEM also highlighted that salt-marsh zonation, based on vegetation assemblages did not influence neither PD nor FD, suggesting that the driving environmental variables are common in both study sites. Environmental filters would then rather act at a landscape scale for both taxa studied (see also Djoudi *et al*., 2018 and Coccia & Farina, 2019). It is therefore also possible that for carabids, the variable influencing both phylogenetic and functional diversity act on a landscape level.

As a conclusion, both spiders and carabids exhibited high correlations between functional and phylogenetic diversities, reinforcing the importance of considering these metrics simultaneously in conservation studies (Webb *et al*., 2002; Swenson, 2011). Interestingly, environmental factors driving functional and phylogenetic diversities differed between taxa, and this, together with percentage of specialist species that differs between the two groups, suggest these two dominant groups of ground-dwelling arthropods differentially react when exposed to stressful factors. Further studies should investigate the role of other, both local (e.g. soil texture) and landscape (e.g. spatial heterogeneity), factors in driving diversity metrics of predatory arthropods in salt marshes. Our study finally highlights that, even in taxa of the same phylum, and occupying the same niche in a highly constrained habitat, functional and phylogenetic diversities can have different drivers, showing different filtering mechanisms and evolutionary history at small spatial and temporal scales.

## Acknowledgements

This work is a contribution to the project PAMPAS ANR-18-CE32-0006 supported by the French National Research Agency. We would like to thank the nature reserve staff of Möeze-Oléron and Lilleau des Nige for field assistance: Fréderic Robin, Jean-Christophe Lemesle, Philippe Delaporte, Vincent Lelong, Julien Gernigon and Lucas Deplaine.

## Data availability statement

Data will be made available on Dryad upon final article acceptance.

**Appendix 1:**
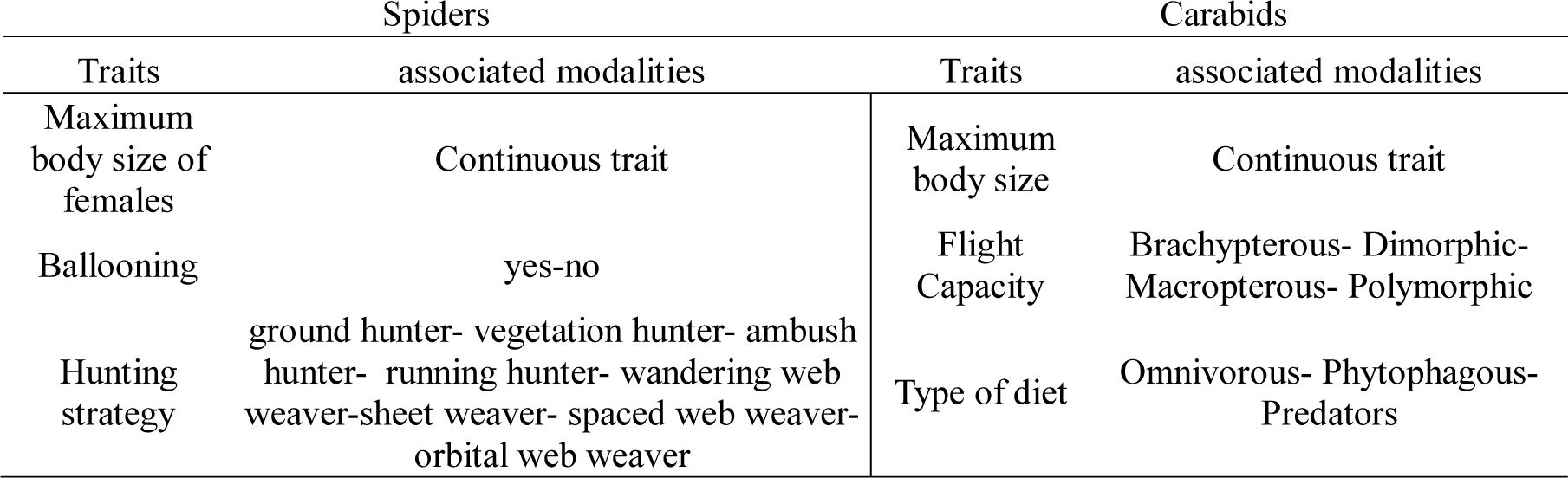
Functional traits used for spiders and associated modalities. The traits were selected according to: Bonte *et al*., (2006) ; Lambeets *et al*., (2008, 2009) ; Albrecht *et al*., (2010) ; Schirmel *et al*., (2012) ; Fischer *et al*., (2013) ; Schirmel & Buchholz, (2013) ; Schirmel *et al*., (2016) ; Gobbi *et al*., (2017) ; Torma *et al*., (2019).

| Spiders |  | Carabids |  |
| --- | --- | --- | --- |
| Traits | associated modalities | Traits | associated modalities |
| Maximum body size of females | Continuous trait | Maximum body size | Continuous trait |
| Ballooning | yes-no | Flight Capacity | Brachypterous- Dimorphic-<br>Macropterous- Polymorphic |
| Hunting strategy | ground hunter- vegetation hunter- ambush<br>hunter- running hunter- wandering web<br>weaver-sheet weaver- spaced web weaver-<br>orbital web weaver | Type of diet | Omnivorous- Phytophagous-<br>Predators |

**Appendix 2:** List of spiders species sampled at site 1 [1] and site 2 [2]. *: halophilic species according to Pétillon *et al*. (2007).

|  |  |
| --- | --- |
| *Agroeca lusatica* (L. Koch, 1875) * | [2] |
| *Agyneta rurestris* (C. L. Koch, 1836) | [1, 2] |
| *Alopecosa albofasciata* (Brullé, 1832) | [1] |
| *Araeoncus humilis* (Blackwall, 1841) | [2] |
| *Arctosa fulvolineata* (Lucas, 1846) * | [1, 2] |
| *Arctosa leopardus* (Sundevall, 1833) | [2] |
| *Argenna subnigra* (O. Pickard-Cambridge, 1861) | [1, 2] |
| *Aulonia albimana* (Walckenaer, 1805) | [1] |
| *Cheiracanthium virescens* (Sundevall, 1833) | [2] |
| *Civizelotes civicus* (Simon, 1878) | [2] |
| *Clubiona neglecta* (O. Pickard-Cambridge, 1862) | [2] |
| *Clubiona stagnatilis* (Kulczyński, 1897) | [2] |
| *Crustulina sticta* (O. Pickard-Cambridge, 1861) | [2] |
| *Diplocephalus graecus* (O. Pickard-Cambridge, 1873) | [1] |
| *Drassodes lapidosus* (Walckenaer, 1802) | [1] |
| *Drassyllus lutetianus* (L. Koch, 1866) | [1, 2] |
| *Drassyllus pusillus* (C. L. Koch, 1833) | [1, 2] |
| *Dysdera crocata* (C. L. Koch, 1838) | [1] |
| *Dysdera erythrina* (Walckenaer, 1802) | [1] |
| *Enoplognatha mordax* (Thorell, 1875) * | [1, 2] |
| *Erigone longipalpis* (Sundevall, 1830) * | [1, 2] |
| *Gnathonarium dentatum* (Wider, 1834) | [2] |
| *Haplodrassus minor* (O. Pickard-Cambridge, 1879) * | [2] |
| *Haplodrassus signifer* (C. L. Koch, 1839) | [1] |
| *Heliophanus flavipes* (Hahn, 1832) | [1] |
| *Heliophanus kochii* (Simon, 1868) | [1] |
| *Leptodrassus albidus* (Simon, 1914) * | [1, 2] |
| *Liocranoeca striata* (Kulczyński, 1882) | [1] |
| *Micaria pulicaria* (Sundevall, 1831) | [2] |
| *Oedothorax apicatus* (Blackwall, 1850) | [2] |
| *Oedothorax fuscus* (Blackwall, 1834) | [2] |
| *Ozyptila simplex* (O. Pickard-Cambridge, 1862) | [2] |
| *Pachygnatha listeri* (Sundevall, 1830) | [2] |
| *Pardosa amentata* (Clerck, 1757) | [1, 2] |
| *Pardosa prativaga* (L. Koch, 1870) | [2] |
| *Pardosa proxima* (C. L. Koch, 1847) | [2] |
| *Pardosa purbeckensis* (F. O. Pickard-Cambridge, 1895) * | [2] |
| *Pardosa vittata* (Keyserling, 1863) | [2] |
| *Phaeocedus braccatus* (L. Koch, 1866) | [1] |
| *Pisaura mirabilis* (Clerck, 1757) | [2] |
| *Pocadicnemis juncea* (Locket & Millidge, 1953) | [1, 2] |
| *Porrhomma microphtalmum* (O. Pickard-Cambridge, 1871) | [1, 2] |
| *Pseudeuophrys obsoleta* (Simon, 1868) | [1] |
| *Pulchellodromus pulchellus* (Lucas, 1846) | [1, 2] |
| *Stemonyphantes lineatus* (Linnaeus, 1758) | [1, 2] |
| *Tenuiphantes tenuis* (Blackwall, 1852) | [1, 2] |
| *Titanoeca hispanica* (Wunderlich, 1995) | [1, 2] |
| *Trachyzelotes fuscipes* (L. Koch, 1866) | [1, 2] |
| *Walckenaeria unicornis* (O. Pickard-Cambridge, 1861) | [1, 2] |
| *Xysticus kochi* (Thorell, 1872) | [1, 2] |
| *Zelotes apricorum* (L. Koch, 1876) | [1] |
| *Zelotes atrocaeruleus* (Simon, 1878) | [1] |
| *Zelotes mundus* (Kulczyński, 1897) * | [2] |
| *Zelotes tenuis* (L. Koch, 1866) | [1, 2] |
| *Zodarion italicum* (Canestrini, 1868) | [1] |

**Appendix 3:**
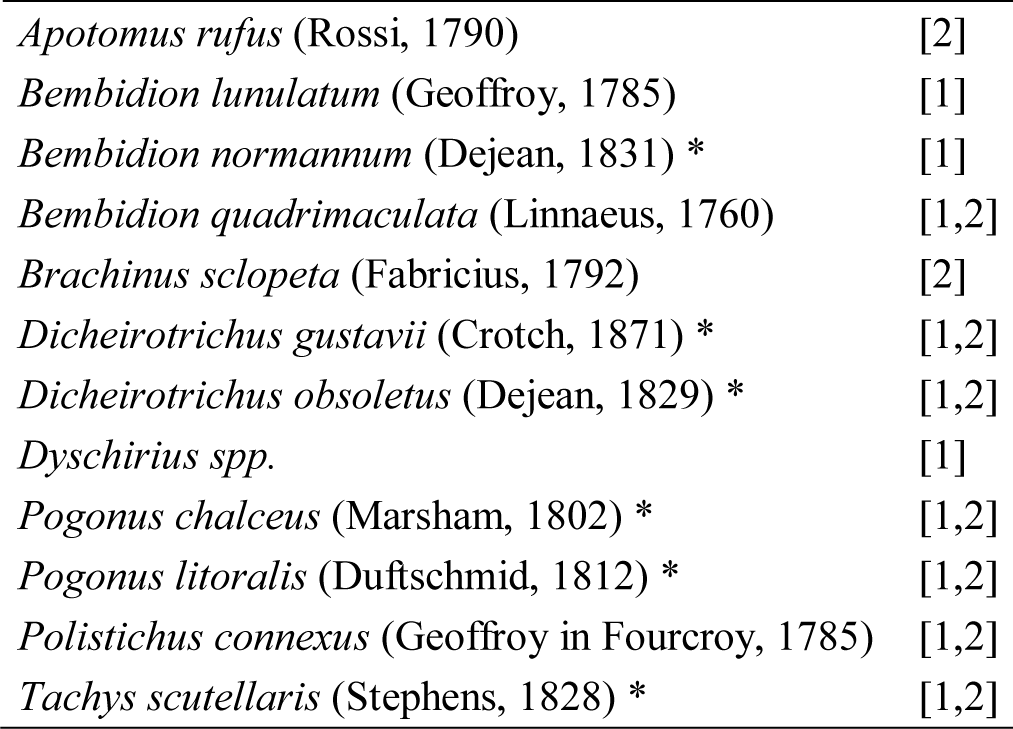
List of carabids species sampled at site 1 [1] and site 2 [2]. The halophilic species are annotated with a star, and considered halophilic according to Pétillon *et al*. (2007) and George *et al*. (2011).

**Appendix 4:**
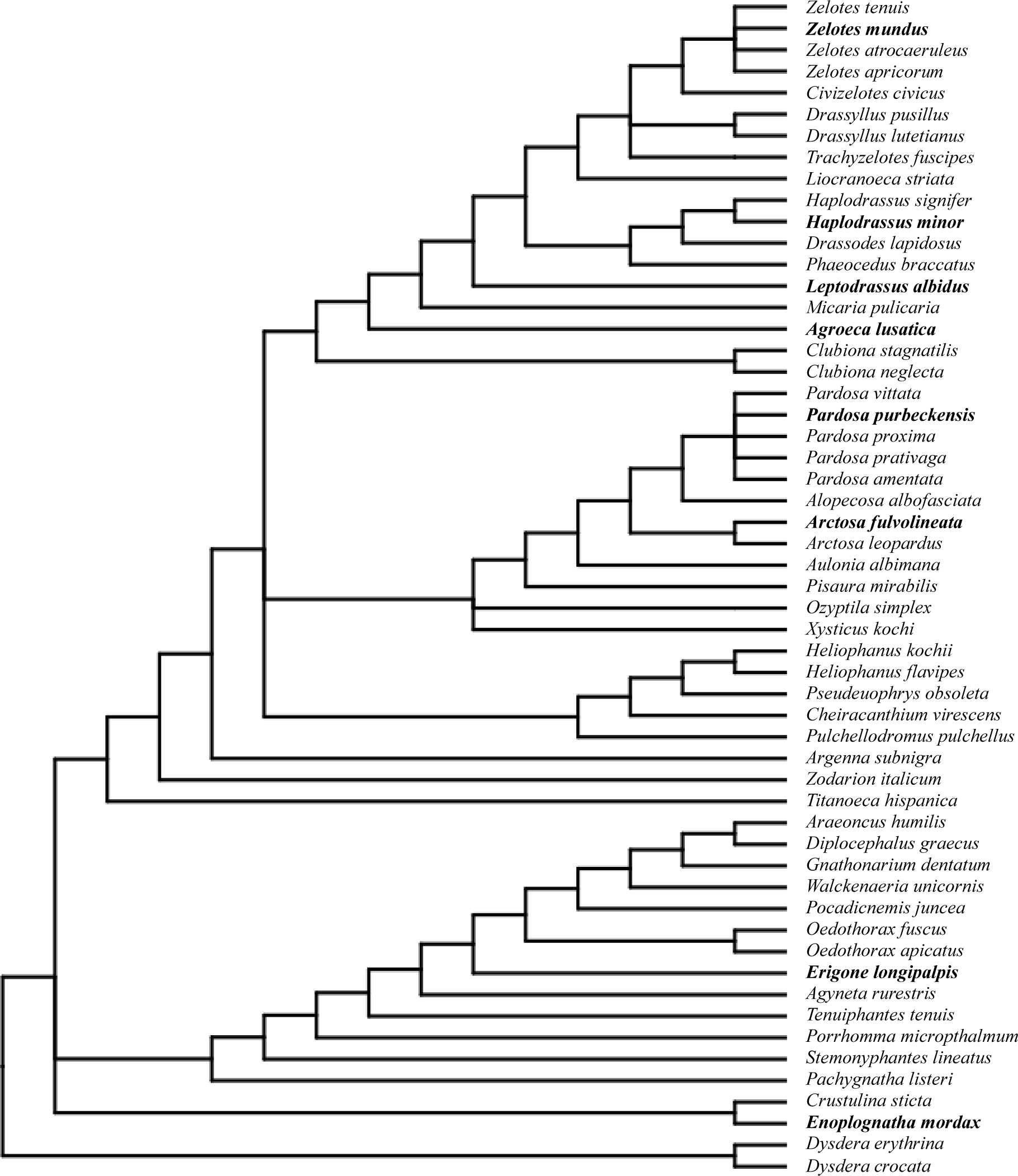
Phylogenetic tree of spider species sampled. Halophilic species are in bold according to Pétillon *et al*. (2007)

**Appendix 5:**
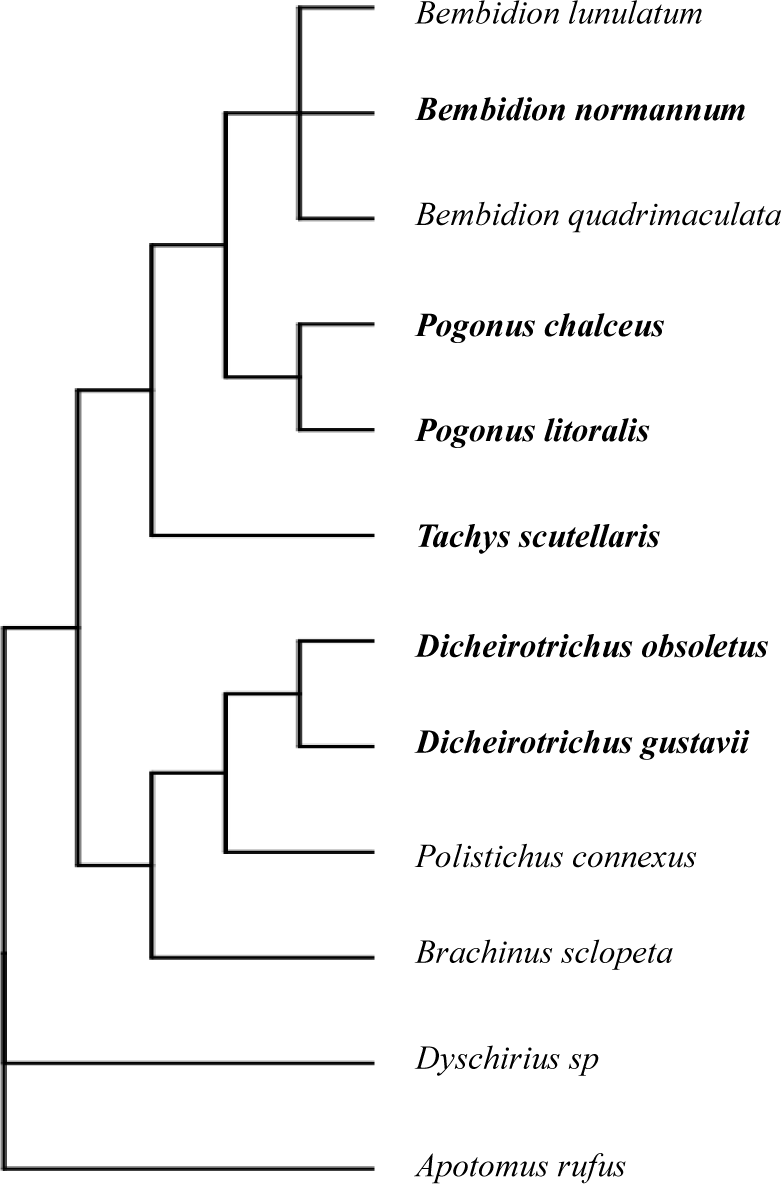
Phylogenetic tree of carabids species sampled. Halophilic species are in bold according to Pétillon *et al*. (2007) and George *et al*. (2011).

